# An Ultra-Stable and Dense Single-Molecule Click Platform for Sensing Protein-DNA Interactions

**DOI:** 10.1101/2020.11.25.397737

**Authors:** Emiel W. A. Visser, Jovana Miladinovic, Joshua N. Milstein

**Affiliations:** Department of Chemical and Physical Sciences, University of Toronto Mississauga, Mississauga, Canada; Eindhoven University of Technology, Eindhoven, The Netherlands; Department of Physics, University of Toronto, Toronto, Canada

**Author notes:** Emiel W.A. Visser – Department of Chemical and Physical Sciences, University of Toronto Mississauga, Mississauga, Canada and Department of Applied Physics, Eindhoven University of Technology, Eindhoven, The Netherlands;, Joshua N. Milstein – Department of Chemical and Physical Sciences, University of Toronto Mississauga, Mississauga, Canada and Department of Physics, University of Toronto, Toronto, Canada.

**Keywords:** TPM, Protein-DNA interactions, High-Stability, Click chemistry, H-NS, Biomarker sensing platform

## Abstract

We demonstrate an ultra-stable, highly dense single-molecule assay ideal for observing protein-DNA interactions. Stable click Tethered Particle Motion (scTPM) leverages next generation click-chemistry to achieve an ultrahigh density of surface tethered reporter particles, has a high antifouling resistance, is stable at elevated temperatures to at least 45 °C, and is compatible with Mg^2+^, an important ionic component of many regulatory protein-DNA interactions. Prepared samples remain stable, with little degradation, for > 6 months in physiological buffers. These improvements enabled us to study previously inaccessible sequence and temperature dependent effects on DNA binding by the bacterial protein H-NS, a global transcriptional regulator found in *E. Coli*. This greatly improved assay can directly be translated to accelerate existing tethered particle based, single-molecule biosensing applications.

## Article text

Dynamic interactions between proteins and DNA are fundamental to the function and regulation of many cellular processes, ranging from DNA organization and gene transcription, to DNA replication and repair. ^1^ A versatile single-molecule technique for studying protein-DNA interactions is Tethered Particle Motion (TPM) in which a reporter particle is bound to a substrate by a DNA tether. ^2,3^ The Brownian motion of the reporter particle is tracked under a microscope while the proteins in solution interact with the DNA. ^4,5^ The Brownian motion reflects changes in the flexibility or conformation (e.g. straightening, looping or kinking) of the DNA tether. Utilizing TPM, protein-DNA interaction, ^6,7^ transcription ^3,8^ and DNA looping ^9,10^ have been successfully studied *in vitro*. Furthermore, significant effort has gone into developing TPM derived sensing techniques to monitor the presence and concentration of medically relevant molecules over extended periods of time, making TPM a promising new method for biosensing applications. Recently, picomolar concentrations of nucleic acids were able to be continuously monitored, over several hours, in blood plasma by Biosensing based on Particle Mobility (BPM), ^11–13^ while fM detection sensitivity to nucleic acids in bodily fluid samples was demonstrated by Single Molecule Tethering (SMOLT). ^14^

The traditional TPM system has a few shortcomings that limit its applicability. The first is that the coupling strategy relies on both a streptavidin-biotin interaction and a much weaker antibody-antigen interaction. The antibody reliant coupling has a limited lifetime (typically a few hours), severely restricting the duration and conditions under which experiments can be performed. Secondly, it proves difficult in practice to achieve a high particle density due to denaturation and random orientation of surface absorbed antibodies. Additionally, the random orientation of the antibodies on the surface adds a variable length up to 15 nm to the tether.

Here we demonstrate a significantly improved TPM assay we dubbed stable click Tethered Particle Motion (scTPM). In scTPM, the traditional antibody coupling strategy is replaced with trans-cyclooctene (TCO) – tetrazine bio-orthogonal click-chemistry at the surface as illustrated in Figure 1a. ^15^ The result is a strong covalent bond, which we find to be stable for months, and thus unlikely to break within the timeframe of a typical experiment or at higher, physiologically relevant temperatures. The TCO activated surface is suitable for the attachment of other molecules of interest, and when combined with streptavidin coated maleimide particles passivated with covalently linked BSA, displays a remarkably high anti-fouling resistance, creating a flexible platform for molecular TPM studies or biosensing applications.

**Figure 1.**
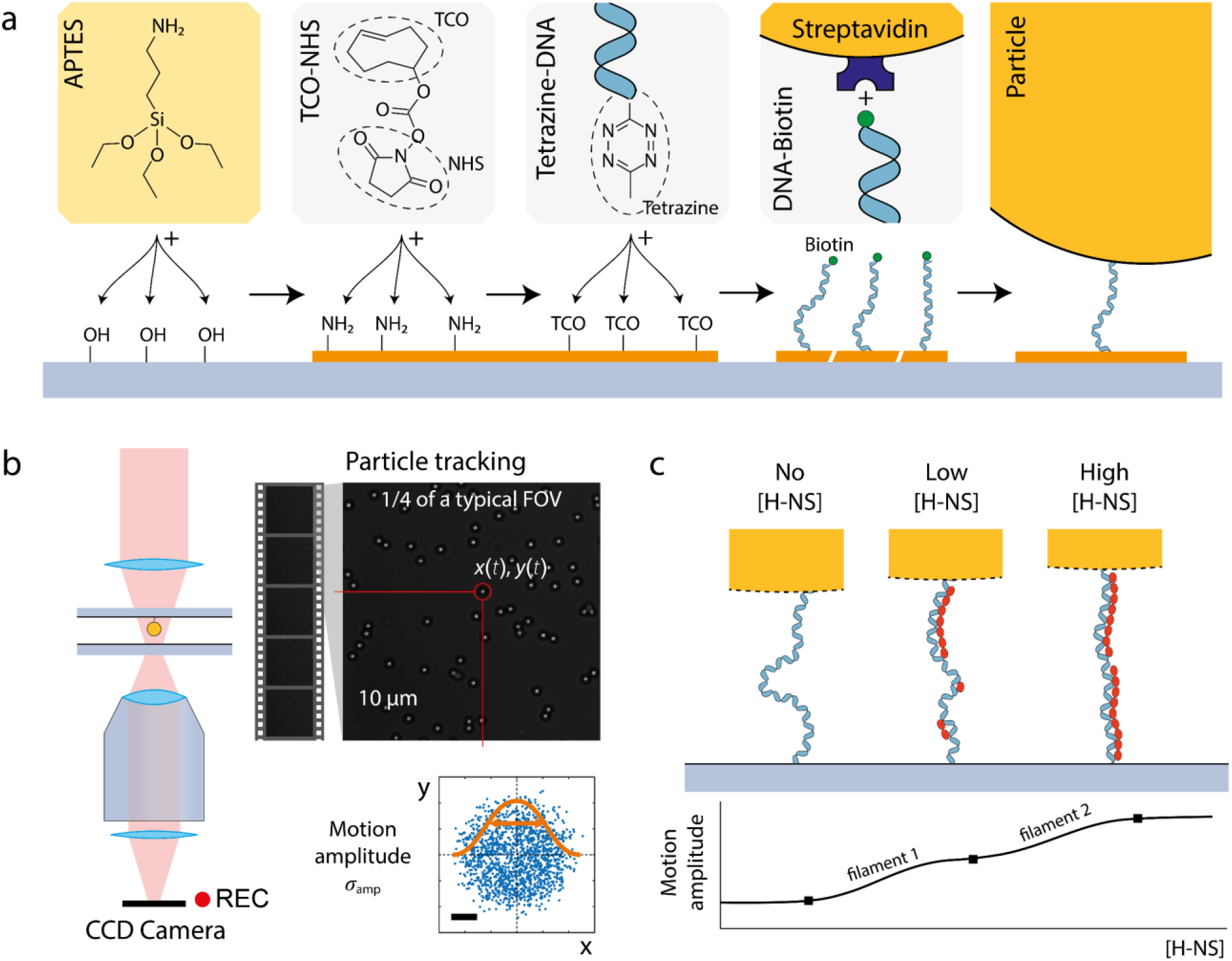
Illustration of the high-stability TPM implementation. a) Preparation steps of the surface to tether particles using the TCO-tetrazine click chemistry. Left-to-right: APTES functionalization of bare glass, attachment of TCO groups using TCO-NHS, surface attachment of tethers using biotin and tetrazine labeled DNA, attachment of streptavidin labeled particles to biotin ligands on the DNA. b) Microscopy setup: the particle motion is recorded using a CCD camera on a brightfield microscope. Video data is analyzed and the motion pattern of the particles determined. c) Illustration of the effect of [H-NS] on the motion amplitude. H-NS binds to DNA and forms filaments in the stiffening mode, increasing the end-to-end distance of the tether. Analysis of the motion amplitude as function of [H-NS] reveals the equilibrium dynamics of the filamentation.

The scTPM system was applied to probe sequence dependent DNA interactions with the Histone-like Nucleoid-structuring protein (H-NS), a highly abundant regulatory protein found in proteobacteria such as *E. coli* and *Salmonella*. ^16–18^ H-NS acts as a global transcriptional regulator and is key to the silencing of many horizontally acquired, xenogeneic genes associated with virulence and drug resistance. ^16^ While H-NS is well studied at both the genomic and ensemble level, the biophysical mechanism it employs to regulate gene expression remains controversial. ^19^ Unlike traditional transcription factors, instead of binding strongly at a certain regulatory sequence, H-NS is thought to loosely associate to AT-rich DNA where it cooperatively forms extended oligomeric filaments (although certain higher affinity sequences are thought to act as nucleation sites for filament formation).

The extreme stability and high-density of scTPM enabled us to acquire the statistics needed to reveal sequence dependent effects on the binding of H-NS to promoter length DNA sequences (see Figure 1c). We then performed protein concentration series to assess mechanical stiffening of the DNA through H-NS binding, for different trial sequences and at elevated temperatures inaccessible to traditional TPM assays.

## Results and discussion

scTPM samples were prepared as shown in Figure 1. A detailed description of the preparation method is given in the Supporting Information. We first verified the tethering quality of scTPM by studying the behavior of the tethered particles as a function of the DNA concentration (a proxy for DNA surface density). The particle motion was analyzed, and selection criteria were set to distinguish between single, multiple tethered and stuck particles (see SI and Figure S2). ^20^ At [DNA] = 0 where particles can only be non-specifically bound to the surface, we typically find about 3 particles (range 0 to 20 particles) per field-of-view. Increasing the DNA surface density leads to an increase in bound particles, with a maximum of single bound particles at 2 pM (118 ± 12 particles per 84.5 × 84.5 μm^2^. This optimum is followed by a decrease in single bound particles and an increase in multiple bound particles and finally to the formation of particles completely stuck through DNA binding. This confirms the full functionality of the scTPM system. We routinely achieve a density of ∼4.2 particles per 100 µm^2^, which approaches the theoretical maximum density of 20 particles per 100 µm^2^ (see SI for details). Imaged on a microscope system with a 10x instead of a 60x objective, we managed to record 5084 properly tethered particles in a single field-of-view (see Figure S6) demonstrating the scalability and high-throughput character of the scTPM assay.

Next, we generated a library of different DNA tether lengths which we incorporated into the scTPM system. The resulting motion amplitudes are shown in Supplementary Figure S3 for the scTPM coupling strategy and compared to simulation results ^20^ and the analytical model described by Segall *et al*. ^21^ An excellent match between experimental and simulation results is found, indicating the proper functionality of the generated tethers and the scTPM system.

Control experiments have been performed using freely diffusing particles in buffer containing up to 5 pM of MgCl_2_ and up to 10 µM H-NS to determine non-specific binding. We typically observed less than 5 particles bound per field-of-view (84.5 × 84.5 μm^2^). The stability of the improved TPM system was assessed through extended observation. Samples were prepared with particles tethered to the surface, closed with tape, stored in the fridge and measured for a period of up to 180 days. A few samples were observed to fail through mechanisms unrelated to the scTPM system (some samples developed a bacterial infection and a few samples dried by evaporation of the buffer through the semipermeable tape used to cover some of the holes). The working samples demonstrate remarkable stability where the number of particles and the observed binding types remained the same after 180 days, as illustrated in Figure S5, indicating that long term experiments can be performed with the scTPM system.

To demonstrate the utility of the scTPM system in biophysical applications, we applied it to study sequence dependent interactions between short, promoter length DNA sequences and H-NS. The influence of sequence on DNA binding by H-NS has proven difficult to study by single-molecule techniques and limited results have been published.^19,22^

Our results are shown in Figure 2. We consider H-NS binding with a synthetic 50% GC control strand with no strong nucleation site in absence of Mg^2+^. With increasing [H-NS] the motion amplitude increases, indicating that H-NS binds to DNA in the filamentous, stiffening mode.^23^ A structured interaction pattern is observed with three rising flanks and two plateau regions around 20 nM and 200 nM H-NS, indicating that a complex interaction between DNA and H-NS exists even without the presence of a strong nucleation site. We hypothesize that this indicates the existence of three separate filamentation events in which segments of the DNA become covered at different [H-NS]. It is likely that low and high affinity regions and nucleation sequences dictate filament formation at different [H-NS] and/or that expanding filaments collide with increasing [H-NS]. Next, we study the effect of GC-content by comparing the 50% GC DNA to AT-rich DNA (42% GC – also no nucleation site). The interaction curve shifts to higher concentrations and the plateaus apparent in the 50% GC DNA construct are no longer apparent, with perhaps just one weaker plateau observed around [H-NS] = 100 nM. The apparent lower affinity is in contrast with the literature where, in general, a stronger affinity is observed between H-NS and AT-rich DNA.^16^ It is likely that this general trend does not hold for all sequences and that sequence specific behavior has a profound effect on H-NS binding.

**Figure 2.**
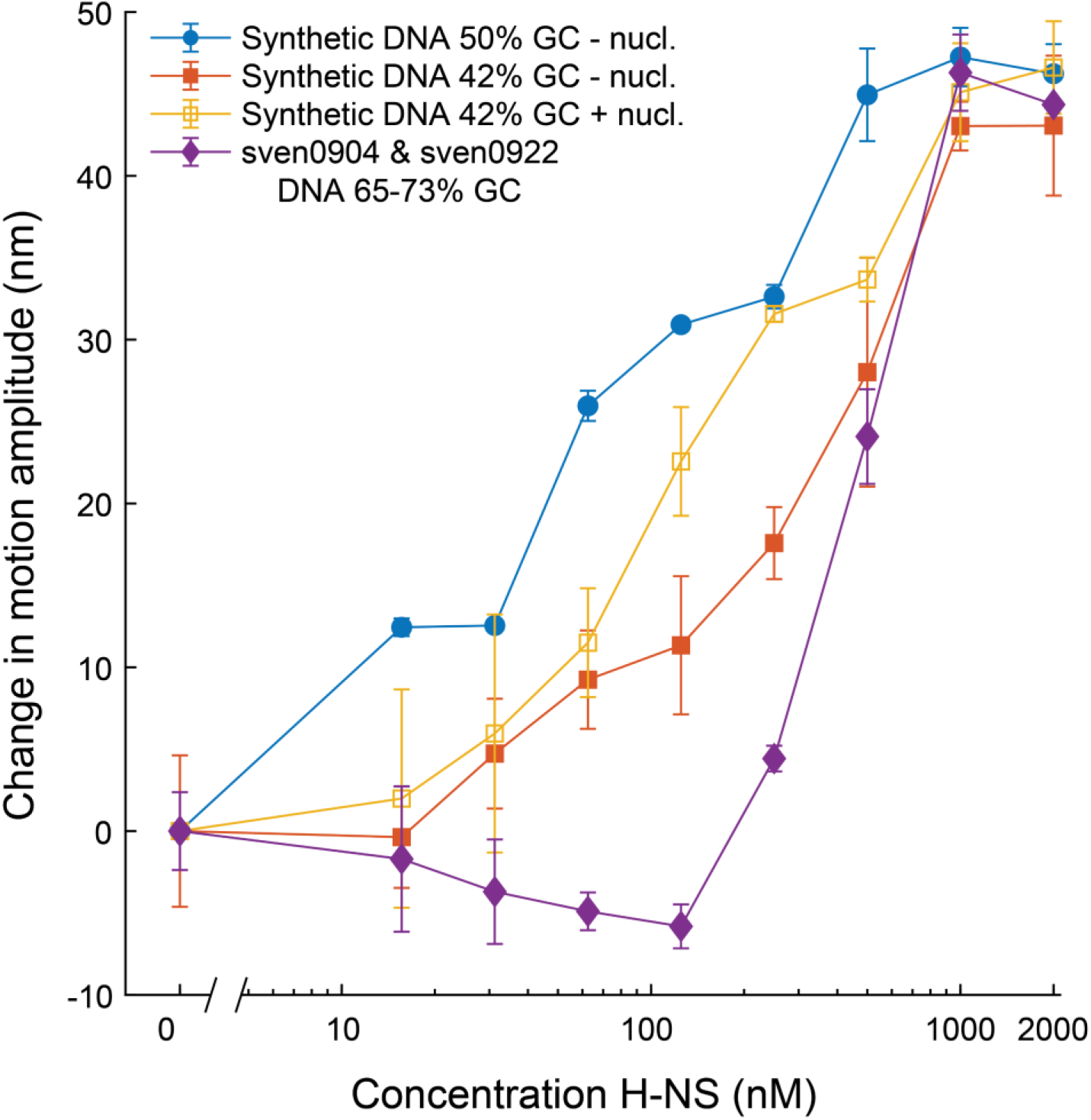
change in (major) motion amplitude A_major_ observed as function of the H-NS concentration for four different DNA tether sequences: 50% GC-content no strong nucleation site (blue circles), 42% GC-content with (open yellow squares) and without (closed red squares) a Pribnow box nucleation site, and natural gene sections sven0904 and 0926 (purple diamonds). Data points represent the average of multiple measurement series, error bars indicate the standard deviation of those values. All curves have been offset such that ΔA_major_ = 0 for [H-NS] = 0 nM.

The effect of a nucleation site was assessed by repeating the experiment with the same 42% GC-content synthetic DNA, but with the bacterial promotor Pribnow box sequence `TATAAT` inserted at the center, a sequence known to strongly interact with H-NS.^22^ While the initial increase in the motion amplitude with [H-NS] remains relatively unaffected, the flank of the curve shifts to a lower [H-NS] of 100 nM, which is where the plateau was observed in the original sequence (absent the high-affinity insert). This observation is congruent with the expected stronger interaction between H-NS and the nucleation site and indicates that the central stepwise increase is likely related to the high-affinity sequence.

Then we consider the interaction between H-NS and segments of the promotor region of the genes 0904 (sven0904) and 0922 (sven0922) found in *Streptomyces Venezuelae* (65-73% GC-content).^24^ Genes of *Streptomyces Venezuelae* are regulated by the H-NS analog Lsr2. Because Lsr2 and H-NS show a remarkable overlap in their interaction with DNA, the interaction of these gene fragments and H-NS is of interest.^22^ The interaction curve in Figure 2 shows an initial slight decrease and subsequent strong rising flank. The shift to higher [H-NS] of the interaction curve agrees with the known lower affinity of H-NS for higher GC-content DNA. We hypothesize that the initial drop may be caused by H-NS inserting, perhaps into TA-steps, in the DNA and using its AT-hook to exacerbate the natural bend in AT. At higher [H-NS], filament formation takes over and stiffening is observed again. To the best of our knowledge such structured, concentration dependent interactions we report here between DNA and H-NS have not been observed before.

Finally, we demonstrate the extended application of the scTPM system by studying H-NS at elevated, biologically relevant temperatures. H-NS has previously been implicated in temperature sensing.^25^ In addition to the usual reduced binding affinity at higher temperatures, H-NS has a thermosensitive switch that blocks DNA binding at elevated temperatures, affecting the regulation of gene expression (*e*.*g*. as bacteria transfer from ambient environments to the temperatures inside the human body).^25^

As a control we observe the effect of temperature on the sven0926 DNA tether (63.1% high GC-content) with temperatures increasing from 22 °C to 45 °C in Figure 3. The control melting curve (H-NS^-^) shows that temperature has no significant effect on the observed motion amplitude. This matches our expectations. In the Worm-Like-Chain description of dsDNA as a flexible polymer the temperature effects on the persistence length of dsDNA^26^ cancel out with the Boltzmann factor. As the temperature has no effect on the observed amplitude, we thus conclude that an observed change in the motion amplitude is likely due to a change in the H-NS filament along the DNA.

**Figure 3.**
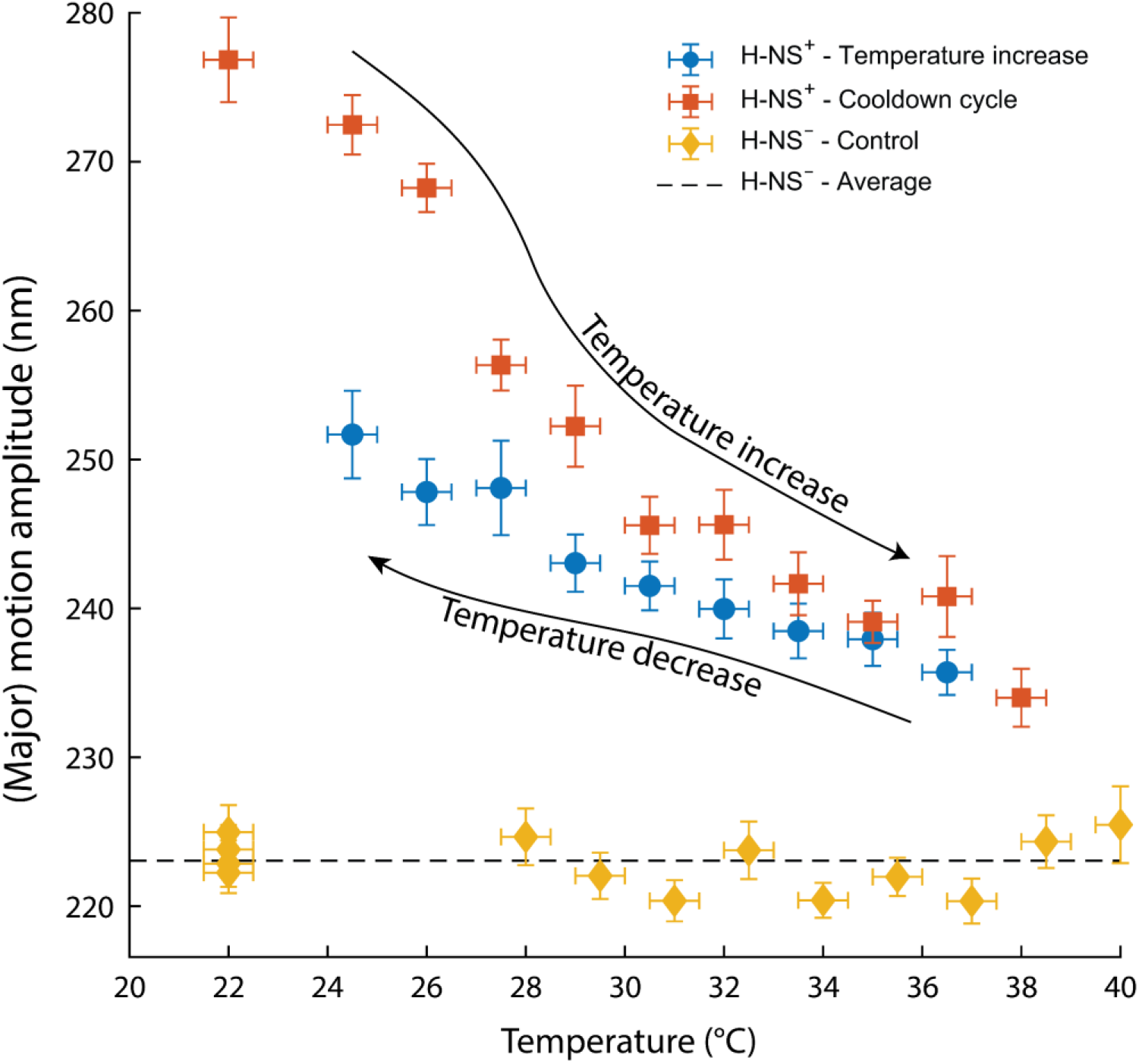
Effect of temperature on the amplitude of motion of 3.5 pM sven0926 tethered particles in the absence of H-NS (H-NS^−^) and in presence of 1 µM H-NS (H-NS^+^). The curves with temperatures increasing and decreasing are shown separately. Horizontal error bars indicate the 0.5 °C uncertainty in the temperature, vertical error bars indicate the standard error of mean of the observed motion amplitude.

We then observed the effect of temperature on DNA interacting with [H-NS] = 1 μM at temperatures increasing from room temperature to 38 °C (1°C above gut temperature) in steps of 1.5 °C. The time between measurements was approximately 15 minutes to allow for temperature equilibration on the setup. The measurements show a decreasing amplitude with increasing temperature, indicating a loss of DNA stiffening by H-NS. This effect could result either from H-NS unbinding from the DNA or from a change in the interaction of H-NS with the DNA and/or neighboring H-NS molecules. The curve shows a gradual decrease in the motion amplitude with the steepest decrease occurring between 26 °C and 27 °C. At 38 °C the curve approaches the bare DNA (H-NS^−^) curve, indicating that only some of the stiffening effect remains. An autoinhibitory temperature switch in H-NS is known to block self-association at elevated temperatures.^25^ Our results indicate that the temperature switch has a continuous response over the entire range of 22 °C to 38 °C.

After the temperature increases, the temperature was decreased to 24 °C in steps of 1.5 °C and the particle motion was recorded at each stop. A partial recovery of the motion amplitude was observed. A full recovery was not observed likely because of denaturation of H-NS over the ∼4.5-hour measurement period. The recovery indicates that the effect is, at least, partly due to a reversible change, as expected for a molecular temperature switch.

## Conclusion

We have demonstrated a greatly improved, high stability TPM system (scTPM) to study interactions between proteins and DNA. Where traditional TPM relies on weak antibody coupling, which dissociates more rapidly at increasing temperatures and adds between 0 and 15 nm of variability to the tether length due to its orientation, the scTPM systems creates strongly bound particles with a well-controlled tether length with no length variation. The ultra-stable click-chemistry of scTPM enables long-term studies, far exceeding the few hours obtainable with traditional TPM (measurement ready samples can remain stable for 6 months or longer), and measurements at elevated temperatures up to at least 45 °C. As far as we are aware, such a long-lived TPM system has not been reported before in the literature. The high reporter particle densities we demonstrate ensure massively parallel experimentation. With a density of ∼4.2 particles per 100 μm^2^, scTPM is on par with a high-throughput TPM assay leveraging surface micropatterning (requiring more complicated microfabrication) to optimize the particle-density to ∼2.7 particles per 100 μm^2^. ^27^ By combining micropatterning and covalent click-chemistry we would be able to approach the theoretical limit of ∼20 particles per 100 μm^2^ for a system with 1 μm particles tethered with 1kb DNA.

These improvements enabled us to study previously inaccessible sequence and temperature dependent effects on the interaction between H-NS and DNA. We find that H-NS interacts with DNA sequences in a highly structured manner, with H-NS filaments likely forming in stages with increasing [H-NS]. The introduction of a 6 bp H-NS nucleation site in the sequence was observed to significantly lower the concentration at which the filament starts to form. Likewise, H-NS filaments along the DNA were observed to soften at increasing temperatures, while regaining their original stiffness upon cooling, hinting at a biophysical mechanism behind H-NS’s role as a temperature dependent molecular switch. These results illustrate the sensitivity of scTPM for finely probing novel protein-DNA interactions.

The scTPM system would clearly enhance current single-molecule force spectroscopy methods such as magnetic or optical tweezers. However, we foresee a range of applications for scTPM well beyond biophysical research. For medical applications, single-molecule biomarker monitoring and detection methods based on TPM have been demonstrated previously. ^11–14^ The high-stability and fouling resistance of the scTPM platform can be used to produce biomarker detection sensors that remain functional for weeks to months at a time and under a range of environmental conditions. The TCO click sensing surface forms a flexible platform for the attachment of nucleotides, proteins and enzymes, and the scTPM surface can be readily functionalized with the required ligands for biomarker capture or the biomarker analog for a competition assay. The click-ready surface enables spatial multiplexing measurements using micropatterning techniques such as microcontact printing or microspotting to generate spatially multiplexed sensor surfaces. Different detection ligands in discrete locations would form a spatially multiplexed sensor akin to a DNA microarray plate in which multiple biomarkers can be simultaneously detected. These properties make scTPM an excellent platform for the creation of medical biomarker sensors. Such sensors require long term stability of the sensing platform and low backgrounds, both of which can be provided by scTPM. Incorporated into a TPM based medical sensor, scTPM would enable one to follow the state of patients in critical conditions by reliably monitoring molecular biomarkers, for up to several months, improving the chance of a positive outcome for these cases. We foresee that scTPM will be applied both as a research tool and a medical technology.

## ASSOCIATED CONTENT

### Supporting Information

Details on the Material and Methods, DNA sequences, verification of scTPM: DNA Concentration Series, DNA tether length, scTPM stability, theoretical maximum particle density and high-density large FOV measurement (PDF)

## AUTHOR INFORMATION

### Author contributions

EV and JNM conceived and designed the methodology, measurement system, and experiments. EV and JM performed the experiments. EV performed the simulations. EV, JM and JNM interpreted the results and wrote the manuscript.

## Acknowledgments

The authors thank Paul Piunno for his helpful discussions on optimizing the APTES functionalization, William Navarre for providing the H-NS proteins, and Marie Elliot and Xiafei Zhang for providing the natural DNA templates from *Streptomyces Venezuelae*.

