## Supporting Information for "An Ultra-Stable and Dense Single-Molecule Click Platform for Sensing Protein-DNA Interactions"

Belonging to the manuscript with the title:

### Methods

**Surface functionalization**

Glass cover slips were plasma cleaned for 15 minutes and APTES (99%, Sigma Aldrich, 440140) functionalized for 1 hour in 1 v/v% APTES in anhydrous toluene. Slides were then thoroughly bathed and rinsed in sequence with methanol, ethanol and twice MilliQ water to wash the slides. Slides were dried using a stream of clean compressed air and then baked overnight at 110 °C to crosslink the APTES layer. Slides were stored in a vacuum chamber until further use.

**Flow chamber assembly**

Ten fluid access ports were drilled in glass cover slides using diamond powder coated drill bits. Flow chambers were cut from 300LSE double-sided adhesive film (3M, Saint Paul, USA) using a plotter cutter (CM350e, Brother). The double-sided adhesive was used to attach the APTES functionalized glass to the microscope slides, forming 5 parallel reaction chambers with 2 fluid access ports each.

**Particle functionalization**

Carboxylic functionalized melamine particles (MF-COOH-AR586, microParticles GmbH, Berlin, Germany) were functionalized with Streptavidin (Sigma Aldrich, 85878) and BSA using an EDC (Sigma Aldrich, E6383) coupling strategy. Each time particles were washed or a buffer exchange was required the particles were centrifuged for 30s 10.000x g centrifugation and the supernatant was removed using a pipette. Particles were incubated 1 hour in 0.01 M NaOH and washed trice with MES buffer (25 mM, pH 4.5). The washed particles were activated in a solution of 10 mg EDC in 150 µL anhydrous DMSO (≥99.9%, 276855, Sigma Aldrich) and 400 µL cold MilliQ on a tube rotator in the fridge. The particles were then washed with cold MilliQ water, sonicated for 10 s and washed with cold MES buffer. Next, they were resuspended in 50 µL Streptavidin in PBS at 0.5 µg/ml for 5 minutes and then mixed with 950 µL 10 mg/ml BSA in PBS and incubated overnight on a tube rotator in the fridge. Particles were then washed twice with 200 µL MES buffer with a 30s sonication in a sonic bath to break up particle clusters. Finally, the particles were resuspended in filter sterilized storage buffer containing 1 mg/ml BSA, 0.05 v/v% Tween-20 in PBS.

**DNA strand assembly**

DNA sequences (linear gBlock Gene Fragments), ssDNA sequences, primers and biotin/azide labeled primers were obtained from IDT (Coralville, IA, USA). The tetrazine labeled primer was prepared by reacting the azide labeled primer 1:1 with methyltetrazine-DBCO (Click Chemistry Tools, Scottsdale, AZ, USA). PCR results were verified on a 1.5% agarose TAE gel, products were cleaned up using PCR cleanup kits (EZ-10 PCR Product Purification Kit, Bio-basic, Amherst, NY, USA) and concentrations measured on a spectrophotometer (Implen Nanophotometer P330). DNA product were stored at -20 °C in TE buffer.

*Synthetic DNA*; the synthetic template DNA was circularized using a ssDNA segment with 18-23 nt overlap at both ends (sequences in SI) in a Gibson assembly reaction using HiFi DNA Assembly Master Mix from New England Biolabs (Ipswich, MA, USA) to generate the template for the next step. The inserted sequence is determined by the ssDNA sequence. The generated template is directly amplified using the biotin and tetrazine labeled primers as the primer binding locations were built into the designed sequence.

*DNA from natural templates*; two consecutive PCR reactions were performed for DNA templates of natural source. The first PCR amplifies the section of interest from the natural template using primers with overhangs to generate the template for the second PCR. The second PCR is then performed with the biotin and the tetrazine labeled primers that attach to the earlier formed overhangs forming the final product with the appropriate labels.

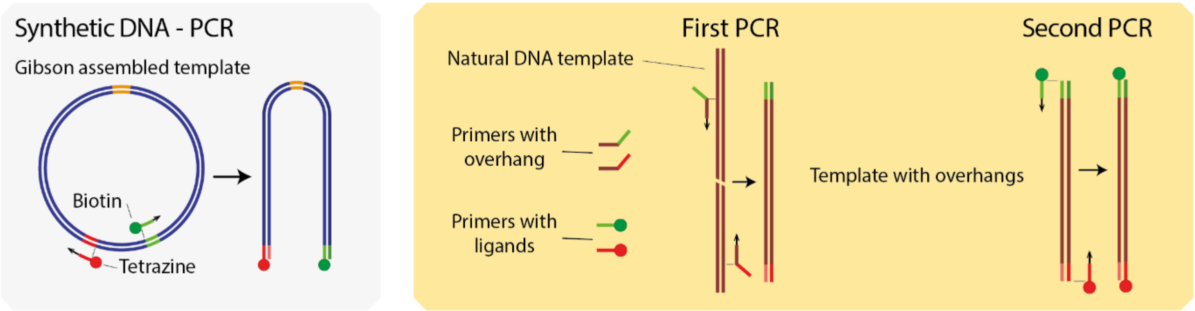


**Figure S1.** *DNA tether production. Synthetic DNA (gray panel): DNA nucleation points were inserted into the tether template using Gibson assembly of a synthetic dsDNA section and a ssDNA insert. From a naturally occurring template (orange panel): a first PCR generates the tether template using two primers with overhangs (green and red). Tether templates are amplified in a final PCR using primers labeled with biotin and tetrazine.*

**TPM system integration**

TCO-PEG4-NHS (Click Chemistry Tools, Scottsdale, AZ, USA) was incubated at a concentration of 1 mM in PBS containing 10 v/v% DMSO (anhydrous, ≥99.9%, 276855, Sigma Aldrich) in the flow chamber for 1 hour at room temperature. Chambers were then flushed with 100 µl MilliQ water, dried using clean air or nitrogen and flushed with 100 µl PBS. DNA tethers were incubated at the required concentration (typically 3.5 pM) for 90 minutes. The chambers were flushed twice with 100 µl PBS. Particles were then flown into the chamber from stock solution at a 15:100 dilution with BSA solution (10 mg/ml BSA in PBS) and incubated for at least 30 minutes. Optionally the chambers were inverted and flushed using the BSA solution to flush out unbound particles.

**Measurements**

H-NS proteins were diluted into HKE reaction buffer (25 mM HEPES, 50 mM KCl, 0.1 mM EDTA) and flown into the chambers through the fluid ports. Measurements were taken on a home build microscope using a Nikon 60x Plan Apo objective (Tokyo, Japan) using brightfield illumination and an Andor iXon Ultra 897 emCCD camera. Typical camera settings: 5 ms exposure time, full frame: 512 x 512 pixels, 25 frames/s, no EM gain. Pixel size calibration (165.5 nm/pixel) was performed through imaging a square grid with known spacing. The data was analyzed using analysis code written in MATLAB described in earlier work. [11]

**Theoretical particle density**

Assuming we are imaging using white light, with an objective with $NA = 1$, assuming $\lambda = 700\ \mathrm{nm}$ as a worst-case estimate, the diffraction limit is $d = 350\ \mathrm{nm}$. With particles $1\ \mu\mathrm{m}$ in size and a maximum excursion of $500\ \mathrm{nm}$ from the center, the particles should be tethered at a distance of 2.35 $\mu\mathrm{m}$ at a minimum to avoid interparticle contact and remain fully optically resolvable. At an organized square grid this leads to a maximum particle density of 18.1 particles per 100 $\mu m^2$, while at a slightly more optimal hexagonal grid this would be a maximum particle density of 22.3 particles per 100 $\mu m^2$. Our TPM system regularly achieves a density near of 4.18 particles per 100 $\mu m^2$ at a random distribution.

**Synthetic DNA sequences**

Underlined: Click and biotin labeled primer sites and Gibson assembly overlap regions **Bold**: Sequence Insert locations (nucleotides are replaced with insert). The sequence shown here is the base sequence without a specific H-NS nucleation site.

**Synthetic tether 42% GC**

Length: 1000 bp. GC-content: 41.6%

**[CLICK chemistry label side]**

AATGGCGGGTCCATGCAGCGTCACAACAATCAGCCCGTTACCCTAATCATTCAGATTTGCGCCCTTGAACGCTACATGTACGAAACCAGCACTAGGT
CAACAATAGGAGTCTTGTAGTTAATATGTAGCCTGGTCGAGTACAGTAGATCCTTCTTACACATTCTATTTATTAAATTAATTCTACAGCAAAACGA
TCATATGCAAATCCACAGTGGCCGATAGACATACATTCACTCCGCTGCTCTGTAATCCTATTTGTATGTTCAGCTAACTTCTACCCATCCCCCGAAA
TTTAGTAGGTTGTGAGATGTCATAGAAGTTCTCGTTCATCCCGTGGGACATCAAGCTTTGCCTTAATAAAGCATTCCGTTCGGGGCAGAAAAAACGC
CTACTGAATTGTGCAATCCTTCTACCTTAACTAAGGTAGCTACCAATATTTAGTTTTTTAGCCTTGCGACAGACTTCCTACTTAGATTGCCACACAT
TGAGC**TAGTGAATCATCCCACACA**ATGGAATGTCCTTAACTCTGGCAGGTAATTAAAGGGAACGTAAAACGCAAAAAAACAGAAAAATAGGCGAATG
AATCTATTTTTACTGTATCGAAGAATGGCTTTGTGGAGGCATGTGTCATGCTAGCGTACAGGGTACTCTAATTATCCATATGGTTCACAAGACACTC
GTTGTTTTCGAATTTACCCTTTATGCGCCGGTTTTCAACCACGCTTATGCTCAACATCGTACCAGACTGATAAGAAATCGGTGTAGCTAAGGACGAA
AGCGACTTTAGGTTCTAACTGTTGACTTTGGCGATAAAGTCAGGAAGCAGACACTGATAGACACGGTTTAGTAGATCGTTTGACGATTAGGTTAAAT
TGAGTGGTTTAATATTGATCTGGCTTTTAGGTGTGTTAATCCGAGTCGAATTAAAAACACCAGTACCCAAAATCAAGCGGGCTCATTACATCGGATA
GTGCAGCTCAGCAGTCTATGAATCCGTAGC

**[BIOTIN label side]**

Used inserts:

| | |
|---|---|
| GC42 Neutral | ACTTAGATTGCCACACATTGAGC**TAGTGAATCATCCCACACA**ATGGAATGTCCTTAACTCTGGCA |
| GC42 Single | ACTTAGATTGCCACACATTGAGC**TAGTGA**TATAAT**CCACACA**ATGGAATGTCCTTAACTCTGGCA |

**Synthetic tether 50% GC**

Length: 1000 bp. GC-content: 49.9%

**[CLICK chemistry label side]**

AATGGCGGGTCCATGCAGCGTCACAACAATCAGCCCTGGCAGGAAATTAAAGGGAACGTACAACGCAAAGAAGCTGGAAAATTGGCGAGGGAATCCT
GTTTCTGTCTATCCAAGAATGGGCATGAGGTGGCAACCGTCGTGCTAGCGTACAGGGTGCACTTTGTAACCATTTGGGACACCGGACACTCGCTGTT
TTCGAAATTACCCTTTAAGCGCGGGTATTGAACCAGGCTTATGCCCAAGATCGTAGCAATACAGAGTTTACCGCATCTTGCCGTAACTGACAAACTG
TGATCCACCACAAGTCAAGCCATTGCCTCTTAGACACGCCGTTAGAGTAATTATGTAAACTTTGCGCGGCTTGACTACGACTCGTTCAGTCACGTCC
GAGGGCACAATCCTATTCCCATTTGTATGTTCAGCTATCTTCTACCCATCCCCGGAAGTTAAGTAGGTCGTGAGATGCCGAGGAGGCTCTCGTTCAT
CCCGTGGGACATCAAATCGTTCAGCGTGGGATTTGCTACAACTCCTGAGCGCTACATGTACGAAACCATGTTATGTATGCACAAGGCCGACAATAGG
ACGTAGCCTTGAAGTTAGTACGTAGCGTGGTCGCATAAGTACAGTAGATCCTCCCCGCGCATCCTATTTATTAAGTTAATTCTACAGCAATACGATC
ATATGCGGATCCGCAGTGGCCGGTAGACACACGTCTACCCCGCTGCTCAATGACCGGGACTAAAGAGGCGACTGCGACGTTCTAAACGTTGGTCCGT
CTGAACCGCCATCCAGGATCACGTCGCCCTGAAAAAAAGATATCAGGAACTCTCCTCCTCAGCAGTCTGGTCTATGGAAACTACAGGACTAACCTTC
CTGGCAACCGGGGGCTGGGAATCTGTCACATGAGTCAAGGTATTTGCTCGATAATCCTCCAGGCATCTAACTTTTCCCACTGCCTTAAGCGGGGATA
GTGCAGCTCAGCAGTCTATGAATCCGTAGC

**[BIOTIN label side]**

**Synthetic tether 59% GC**

Length: 1000 bp. GC-content: 58.6%

**[CLICK chemistry label side]**

AATGGCGGGTCCATGCAGCGTCACAACAATCAGCCGGACTTCTTATTAATTCTTTCATCGTGGGGAGCAGCGGATCTTAATGGATGGCCGCAGGCGG
TGTGGAAGCTAACAGCGCGGGTGGGAGGGTAATCAGCCGTGTCCACCTACACAACGCTAACGGGCGGTTCGGATTCCGCATTGCGCCTACCGGGTGC
CTCAACGGTATCCGCGACTTGCGAAGTGCCTGCTATCCTTGAACGCATACCTCGCCAGCGCGCCGCACTTATTGCGTGAGTAGGGTCGACCAAGAGC
CGCTAGATGCGTCGCTGTGCAAATAGTTGCCGACAGACCGTCGAGTTTAGAGAACGGTGCCAGCATTTTCGGGGGATCTCAATCAAGTATGGGTTGC
GGTGTCTGCACTGTCCCGCGGCTACCCATGGCCTGAAACCCAGCTCGTGTCAAGCCGTCGCCTCTCCGGGACGCCGCGCGAAGTGACACATACACCT
TGCACGGGTTCACCGTAACATATTCTTACGTGTGACGCAGCTGTGTATTCTGCAGAGGTGGCGAACGGGTTCGACACTTCACGGATGGTGGGGATCC
GGGCAAAGGGCGTGTGATTGCGGCCCAACACAGGCGTAGACTACGACGGCGCCGGCTCAGTCGCAGCTCGTGCGCCGTGAATAACGTACTCATCCCA
ACTGATTCTCGGCAGTCTACGGAGCGACACGATTATCAACGGCTGTCTAGCAGTTCTAATCTCTCGCCACCGCCGAAAGCATAAACGACGGGCAGGT
ACGAGAGTGCTAGAACTGGACGTGCCGTTCCTCTGCGAAGAACACCTCGGGCTGTGGCGTTGTTGCGCTGCCTAGATGCAGTGTCGCTCGTATCGCA

CTCGCCTCAACGGCTGCCGCCTTCGCTGCGTCCCTAGACACCCAGCAGTAAGCGCCTTTCGTAGGCGGGGGCACCCCCTGTCAGTGACTGCGGGATA
GTGCAGCTCAGCAGTCTATGAATCCGTAGC

**[BIOTIN label side]**

### DNA from natural templates

Primers for PCR cycle 1:

Forward primer with sticky end: AATGGCGGGTCCATG**CAGCGTCACAACAATCAGCC**

Reverse primer with sticky end: GCTACGGATTCATAG**ACTGCTGAGCTGCACTATCC**

Primers for PCR cycle 2:

Click primer: /Azide/AATGGCGGGTCCATG**CAGC**

Biotin primer: /Biotin/GCTACGGATTCATAG**ACTGCTG**

The click-primer is further reacted with Methyltetrazine-DBCO to generate tetrazine labels on the primer. The sequences of the tethers are shown below. Underlined and **Bold**: sequences added to facilitate biotin and click-chemistry labeling.

#### sven0904

Length: 1094 bp. GC-content: 64.7%

AATGGCGGGTCCATG**CAGCGTCACAACAATCAGCC**CTGCAGCCCTTGTGTCCATATCCCTTGTGACCGGATGCAGTGTCTTCTCGGACAGTGATTCC
GCCGGGGACCAGCGAATCGTCGTCGGTACGACGAGTTCTCCCACCACGCTTGATCCCGCTGCGGCCTGGGACAATTCATGGGAGCTCTTCCGGAACG
TCTTCCAGACCCTGGTGAGTTTCCCGACGGGCAGTACGACTCCGGATTCGGACGCCGCCGACTGCAAGTTCTCGGACACGTCGAGCCGGGTCTTCGA
GTGCAAGCTCGTGGAGGGCCTCACCTTCTCCAACGGGCACAAACTGGACGCCAAGGCCGTGCAGTACTCGATCGAGCGCATCCGCACGATCAACCAC
AAGGGCGGCCCCAACGGCATGCTCGGCTCGCTCGACAAGATCGAGACCGTCGGCGACCGGACCGTCGTCTTCCGGCTCAACAAGTCGGACGCGACCT
TCCCGTTCGTCCTCGCCACCCCCGCGATGTCGCTGGTGGACCCCGCCGAGTATCCGGCGGACCGGCTCCGCACCGACGGCAAGGTCACCGGCTCCGG
GCCGTACGTCCTCGACAAGTACACCGAGCGCGAGACGGCCGAGCTGACCAAGAACCCCACCTACAAGGGCTTCGCGGACCGCAAGAACGGCGGAGTC
ACCATCCGCTACTTCGACGAGTCCGACGCCATGGTCGCCGCGCTCAAGAAGCAGGAGATCGACGCCACCTACCGCGGCCTCACCGCCGAGGAGGTCG
TCGCACTCCAGGACGACAAGCCCGAGAACAAGGGCCTCCAGATCGTCGAGTCGACCGGCGCCGACATCCGCTACCTCGTCTTCAACGCCCAGGACCC
GACCGTCGCCAAGCCGGCCGTCCGCCGGGCCGTCGCCCAGCTCGTCAACCGCGACGAACTGGTCAACAAGGTCTACCAGGGCACCGCCGAGCCCCTG
TACTCGATGGTGCCCAAGGGCATCGCCGCCCACACCACCAAGTTCTTCGACCGCTACGGCTCGCCGAACGCCCAGAAGGCCAAGAAGAT**GGATAGTG**
**CAGCTCAGCAGT**CTATGAATCCGTAGC

#### sven0922

Length: 1133 bp. GC-content: 72.8%

AATGGCGGGTCCATG**CAGCGTCACAACAATCAGCC**CCTGTTGACCGACCTCTTCCGGCGCCACCGCGGACGGACCGCGCTGCGGACCGCCGGACGGA
CCTGGACCTACGAGGAGCTCGACCGGGTCACCAGCGCGCTCGCCCGCCGGATCGACGCCGAGTGCCCCGCGGGCCGCCGCGTCCTGGTCGCCGGGGA
GCACACGGCGGAAGCCGTGGTCTGGGCCCTCGCGGCGATGCGCAGCCACGCCGTGCACACCCCGATGAACCCGGGCCTGCCCGCCGACCGGTTCGAG
GAGTTCGCGCGGGTCGCGGACGCGGCGCTGCTCGTCTGCTTCGAGCGCGAGGCGCTGGTGCGCGGCGAGAAGGCCGGCCTGCGCGCCCTGTACGCCG
GTGATGTCGGCTGGCCCACCGACCCGGCCCCGGCGCCGGCGGACGGCACGGCCGACGAGCCGGCCCGTTCCCGCGTCGCGTACTCCATCTTCACCTC
GGGTTCGACCGGCGACCCCAAGCTCGTCGACGTCGGGCACGGCGGCCTGCTCAACCTCTGCCGCTCCCTGCGGAGGCTGCTCGACATCACCCCCGAC
GACCAGGTCCTGCACTTCGCGTCGCTGTCGTTCGACGCCTCCGTCAGCGAGATCCTCGGCACCCTGTACGCCGGGGCGACCCTCGTCGTCCCCGTGC
GCGACCAGGCCTCCTGGCTGGGGTCCGTCTCGCGGCACCTCGCCGCCCACGGCTGTGACCTCGCCATGCTGTCGCCGTCCGTCTACGCCCGGCTCGA
CGAGGCGGCCCGCAGCCGCATCCGGAAGGTGGAGTTCTGCGGCGAGGCCCTCGGCGAGGGCGAGTACGACAAGGCCGCCCGGTACAGCAGGGTGTTC
AACGCGTACGGCCCGACCGAAGCCACCGTCTGCTTCTCCCTGGCCGAGCTGACCTCGTACACGCCGAGCATCGGCACCCCCGTCGACGGCTTCCGGG
CCTACGTCCGCGACCCCGACTCCGGTGACCACGCCACGGCGGGCACCGGCGAACTCGTCATCGTCGGCGACGGCGTGGCACTCGGCTACGCCGGCGG
CAGCCCGGCCGAGAACGAGGTGTTCGGTACG**GGATAGTGCAGCTCAGCAGT**CTATGAATCCGTAGC

#### sven0926

Length: 1033 bp. GC-content: 62.8%

AATGGCGGGTCCATG**CAGCGTCACAACAATCAGCC**CGACCTGTGTCTCTACGGTGACTCCGGCGTGTTCGAGTCCATGTACCTGGTGGCCTTCCTCT
CCCTGGTCGAGGAGAACATCGAGGACGAGTTCGGCACCGAGGTGTCGCTCACGTCCGAGAAGGCGGTGTCCCGCCGGGTGAGTCCGTTCAGCAGCGT
GCGCAGGCTGATCGGCTTCATCGAGGAAGAGCTGGAACTCGTCGGCGTGCGCGGTTAGCGCAGGTCACCGCCGCCATGATCCTCATGATCACCGGCA
CCCGGACCGGCATCGGCAAGGCCCTCGCGGAGCATTTCCTGGAGGCCGGGCACGACGTCGTCGGGTGCAGCAGAAGGCCCCCGACCATCGACCACCC

GCGCTACCGGCACCACGAGGCGGATCTCGCGGACCCCGGTCAGACGAAACGCATGTTCCGGAGCGTCCGGGCGGAGTACGGAAAGCTCGACGCCCTG
GTGAACAACGCCGGCACCTCGTCGATGAACCATTTCATGACGACCCCGGACGAGGTTTCCCGGAAGATATTCGACATCAATTTCTTCGCCGTGCTGA
ACTGCTGCCGGGAAGCGGTCAAACTGCTGCGGAAATCCACTGAGCAGAGTTCCGCGATCCTGAACGTCTCCACGGTCGCGGTGCCCTGGGCCATCGA
AGGACAGCTCGCCTATTCGGCCAGCAAGAGCGCGGTCGAACAACTCGTCCGCGTGATGAGCAAGGAGTTGTCCACGTTCGGGATCAGGGTGAACGGA
ATCGGACTTCCGCCCGTTCACACCGTATTGACGAGGACCGTTCCCCGGCCAAAGATCGACGCTCTGATCGCACGCCAGGCGATATCCCGTCAGTGCA
CGACGGCGGACATCGTCGGCCCCGTGGAGTTCCTGATCGGCCGGCAGTCGGAGTTCATCACGGGCGAGACGCTCTTTCTAGGGGGTGTGCACTGATG
CGCGACGAACTGCTGCGAAGGTTCGAGG**GGATAGTGCAGCTCAGCAGT**CTATGAATCCGTAGC

**DNA concentration series**

Tethered particle samples were prepared with the sven0926 tether and a tether concentration from 256 pM down to 0.0125 through serial dilution by a factor of 2 with a negative control 0 pM. The particle motion was recorded for 5 fields-of-view per concentration and the particle motion was analyzed. Particles were classified based on the motion parameters. Single tethered particles were defined as having $A_{minor} \in [140,250]$ and $Sym \in [0.75,1]$. Multiple tethered particles: $A_{minor} \in [30,140]$ and $Sym \in [0.2,1]$. Stuck particles: $A_{minor} \in [0,30]$ and $Sym \in [0.2,1]$. The number of particles with a certain classification is shown as function of the tether concentration in Figure S2.

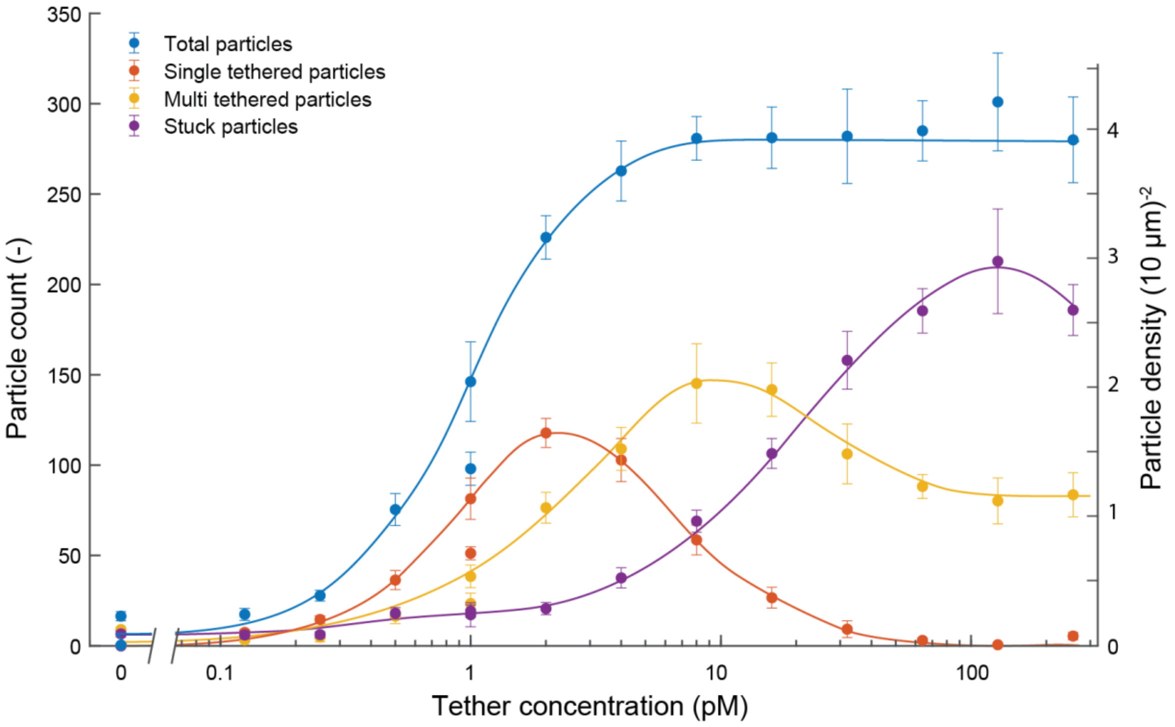


**Figure S2.** *Number of particles classified as single tethered, multi tethered and stuck particles as function of the sven0926 tether concentration. Two separate series are shown, one from 1 to 256 pM and one from 0.125 to 1 pM, both with a 0 pM negative control. Error bars indicate standard deviation between 5 field-of-views. Lines have been drawn in as guides to the eyes.*

Figure S2 shows a monotonous rise in the number of particles with the tether concentration, which is consistent with an increased number of available binding location for particles on the surface. The system saturates at about 280 particles per field-of-view at 8 pM, at which we most likely bind all functional particles. In the control at 0 pM tethers we observed some sample to sample variation between the number of (non-specifically bound particles) one sample showed < 1 particles per field-of-view, while another sample showed about 20 particles per field-of-view. In our experience we typically observe 2 – 4 non-specifically bound particles per field-of-view (data not shown) in our controls. With increasing [DNA] particles become more likely to be surface bound by multiple DNA strands which is observed as that particles with increasing [DNA] first become surface bound with a single tether, then with multiple tethers and finally become stuck. At a DNA concentration above 64 pM the number of particles bound in the respective categories does not change much anymore. This is explained by either the surface TCO groups being saturated, or the streptavidin binding sites on the particles being saturated.

At the optimal concentration in this experiment for single tethered particles (2 pM) a total of 118 ± 12 particles are bound with a single tether. This may be further improved upon by optimizing the density of surface TCO groups and particle streptavidin groups to lower the chance of multiple tether formation.

**DNA tether length**

DNA tethers of different lengths were generated through the PCR based method described in the methods. The selection of single tethered particles was made using selection criteria described in Table S1.

*Table S1: Overview of the tether lengths and selection criteria used to select for single tethered particles.*

| | | Selection criteria | | |
|---|---|---|---|---|
| | Tether length | $A_{minor}$ (nm) | $Sym$ | Tether concentration |
| Sven0926 | 1133 bp | ∈ [140,250] | ∈ [0.75,1] | 1, 2, 4 & 8 pM |
| λ-DNA Primer 2 | 1917 bp | ∈ [200,300] | ∈ [0.75,1] | 1 & 5 pM |
| λ-DNA Primer 7 | 1278 bp | ∈ [175,250] | ∈ [0.75,1] | 5 pM |
| λ-DNA Primer 10 | 3186 bp | ∈ [250,350] | ∈ [0.75,1] | 0.25 & 1 pM |
| λ-DNA Primer 12 | 762 bp | ∈ [150,200] | ∈ [0.75,1] | 5 pM |
| λ-DNA Primer 14 | 4562 bp | ∈ [250,400] | ∈ [0.75,1] | 0.25 & 1 pM |

The selection criteria were based of off the motion distribution plot where the criteria were set to encompass the full population with high symmetry and high amplitude of motion.[28] The $A_{minor}$ and $A_{major}$ was determined from the selected (single) tethered particles. Monte Carlo simulations[20] were performed with a persistence length of 50 nm for dsDNA and a contour length of 0.34 nm per base pair and the *xy* motion pattern coordinates were analyzed as if they are experimental data. The results are shown in Figure S3.

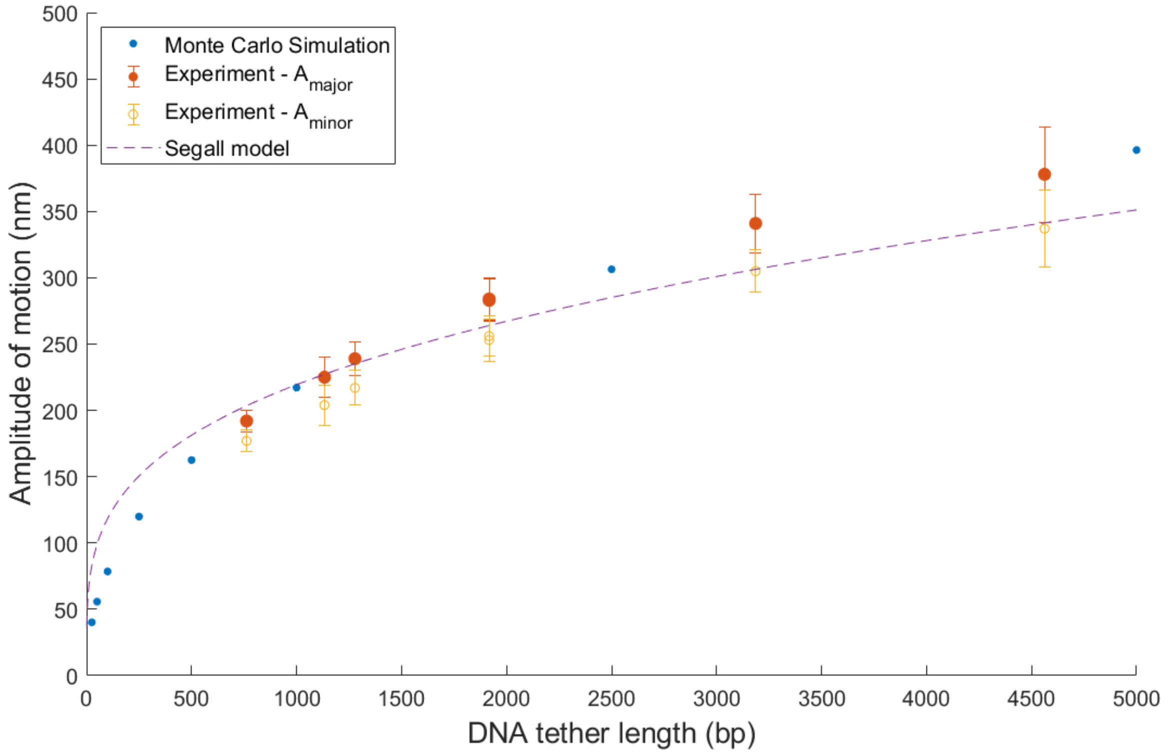


***Figure S3.*** *Amplitude of motion (both $A_{minor}$ and $A_{major}$) for tethers of different lengths compared with Monte Carlo simulation results and the analytical model by Segall et al. shown here as $(\langle r_{\perp}^{2}\rangle/2)^{1/2}$ to be comparable to our definition of the motion amplitude $\sigma_x = \langle x^2\rangle^{1/2}$.[21] Error bars indicate the standard deviation within the observed experimental population. Simulation results have standard deviation < 1 nm and are therefore not shown.*

Figure S3 shows a clear agreement between the simulation results and the experimentally observed major motion amplitude $A_{major}$. Tether lengths were confirmed by gel electrophoresis shown in figure S4. The $A_{minor}$ values naturally lie below $A_{major}$. The difference between $A_{major}$ and $A_{minor}$ is due to the limited sampling time of the particle motion and small imperfections in the tethered particle system.

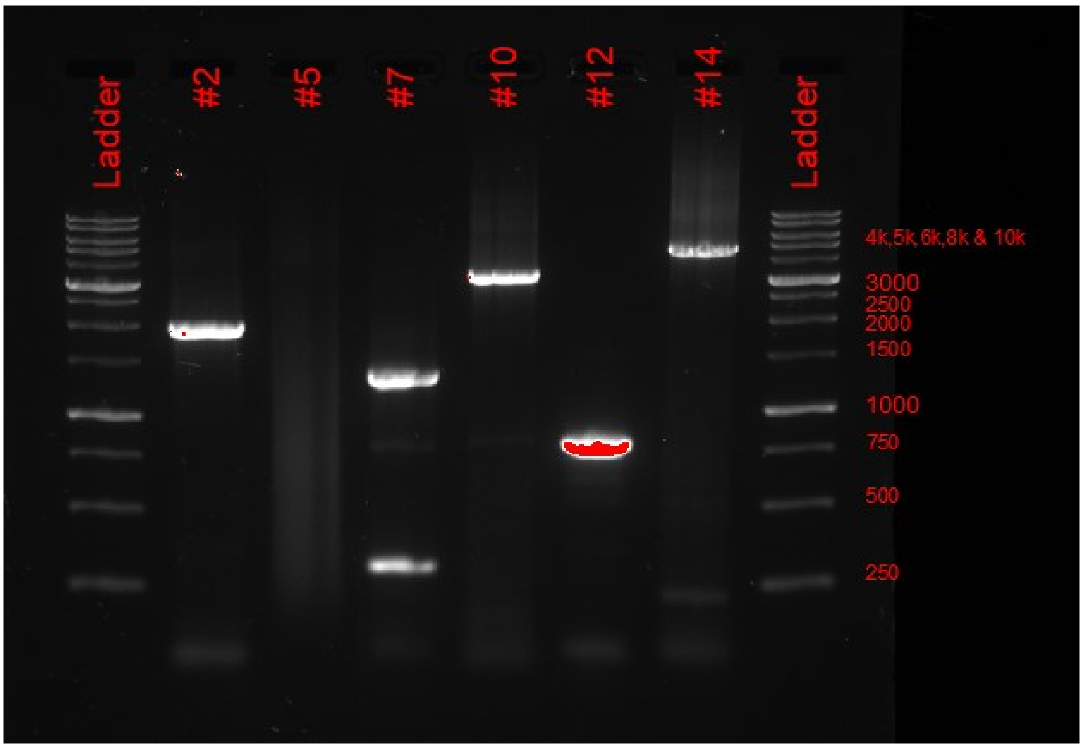


***Figure S4.*** *Gel electrophoresis result of the tethers generated from λ-DNA. The top band was extracted in the lane with primer #7.*

**scTPM stability**

Sample stability was assessed by comparing the particle motion of the same scTPM sample after 7 days and after 180 days. The result is shown in Figure S5.

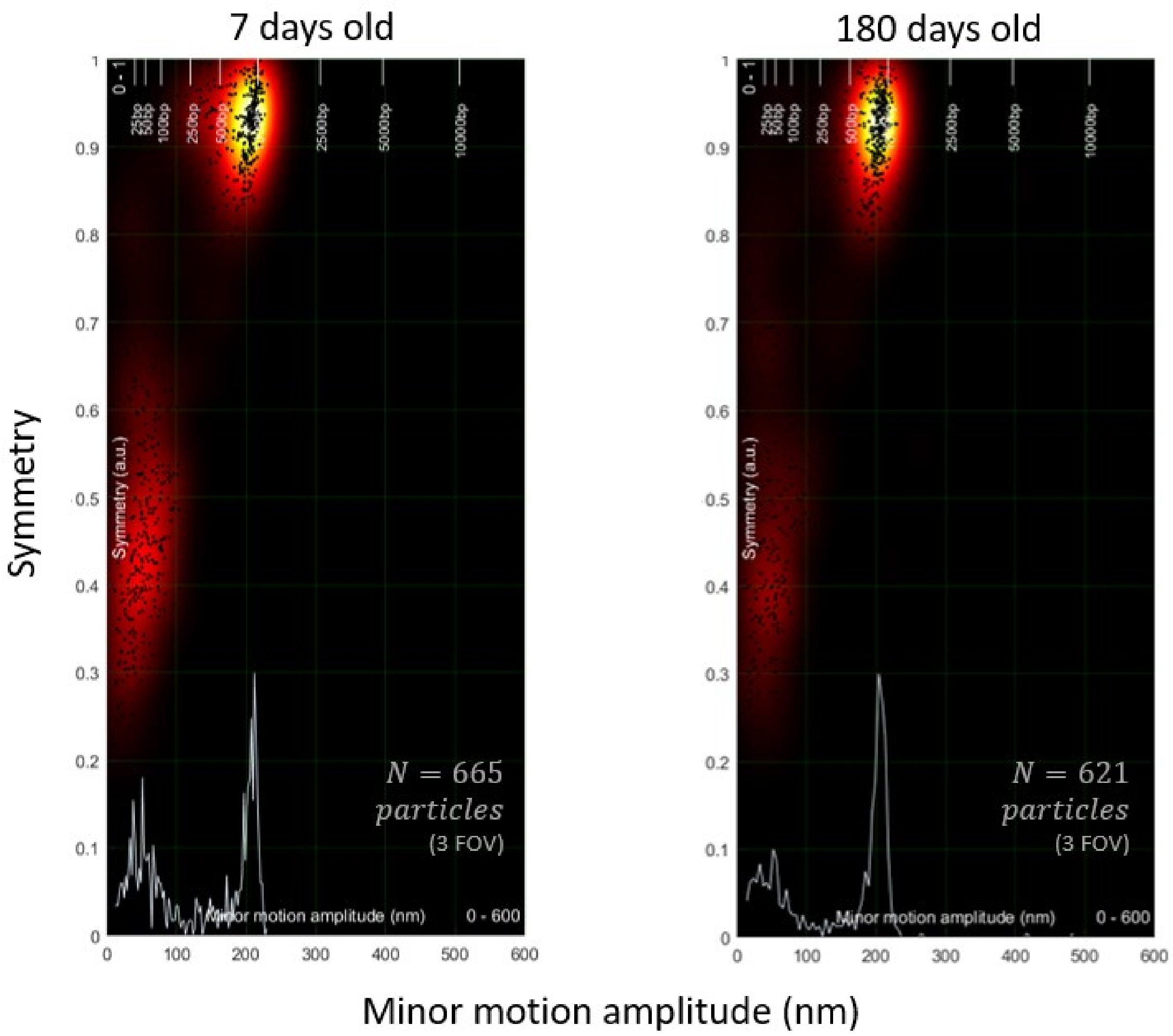


***Figure S5.*** *Comparison between the particle motion distribution of a sample after 7 and 180 days of storage under physiological buffer conditions (PBS, pH 7.4 + 1 wt% BSA) at 4°C. Vertical axes represent the minor motion amplitude* $A_{minor}$ *while the vertical axes represent the symmetry* $Sym$ *of the motion pattern. Each black dot represents a single particle motion. The color map indicates the local density of motion patterns with similar* $A_{minor}$ *and* $Sym$*. The inserted histogram indicates the distribution along* $A_{minor}$*. The top axis shows the expected motion amplitudes for different tether lengths.*

**Wide FOV measurement of tethered particles**

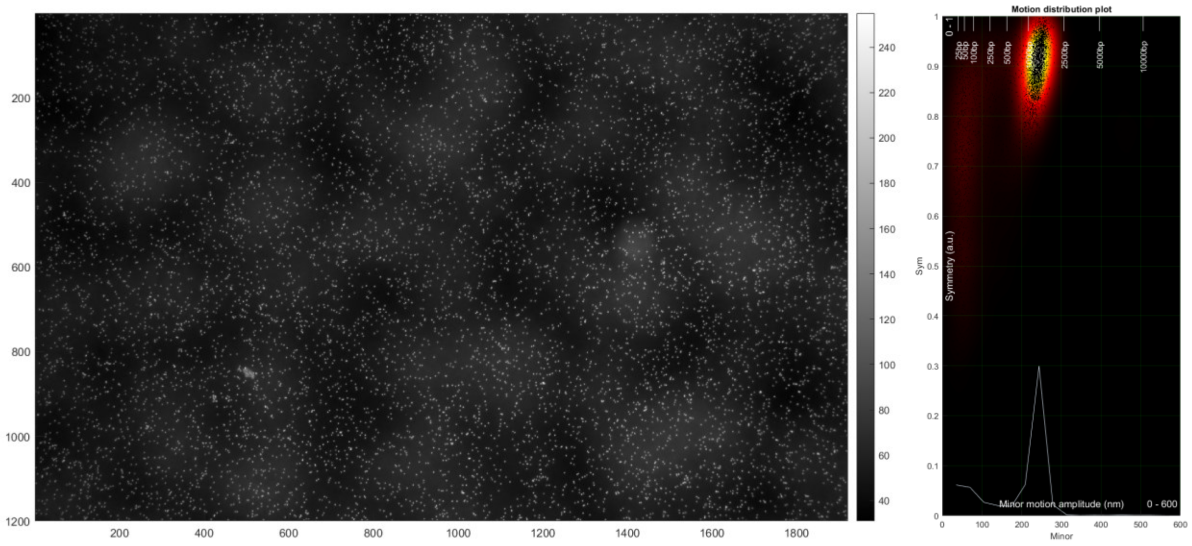


***Figure S6.*** *(left) One FOV (approximately 700 x 1130 µm) at a magnification of 10x. The cloudy background originates from out of focus sedimented particles. (right) A total of 8556 particles were successfully tracked in this particular FOV with 5084 particles exhibiting a motion that corresponds to single tethered particles.*
